# A comparison between full-length 16S rRNA Oxford Nanopore sequencing and Illumina V3-V4 16S rRNA sequencing in head and neck cancer tissues

**DOI:** 10.1101/2024.03.08.584026

**Authors:** Kenny Yeo, James Connell, George Bouras, Eric Smith, William Murphy, John-Charles Hodge, Suren Krishnan, Peter-John Wormald, Rowan Valentine, Alkis James Psaltis, Sarah Vreugde, Kevin Aaron Fenix

## Abstract

**Introduction:** Describing the microbial community within the tumour has been a key aspect in understanding the pathophysiology of the tumour microenvironment. In head and neck cancer (HNC), most studies on tissue samples have only performed 16S ribosomal RNA (rRNA) short-read sequencing (SRS) on V3-V5 region. SRS is mostly limited to genus level identification. In this study, we compared full-length 16S rRNA long-read sequencing (FL-ONT) from Oxford Nanopore Technology (ONT) to V3-V4 Illumina SRS (V3V4-Illumina). To date, this is the largest study using HNC tissues samples to perform FL-ONT of the 16S rRNA using ONT.

**Methods:** Sequencing of the full-length and the V3-V4 16S rRNA region was conducted on tumour samples from 26 HNC patients, using ONT and Illumina technologies respectively. Paired sample analysis was applied to compare differences in diversities and abundance of microbial communities. Further validation was also performed using culture-based methods in 16 bacterial isolates obtained from 4 patients using MALDI-TOF MS.

**Results:** We observed similar alpha diversity indexes between FL-ONT and V3V4-Illumina technologies. However, beta-diversity was significantly different between techniques (PERMANOVA - R^2^ = 0.083, p < 0.0001). At higher taxonomic levels (Phylum to Family), all metrics were more similar among sequencing techniques, while lower taxonomy displayed more discrepancies. At higher taxonomic levels, correlation in microbial abundance from FL-ONT and V3V4-Illumina were higher, while this correlation decreased at lower levels. Finally, FL-ONT was able to identify more isolates at the species level that were identified using MALDI-TOF MS (81.3% v.s. 62.5%).

**Conclusions:** FL-ONT was able to identify lower taxonomic levels at a better resolution as compared to V3V4-Illumina 16S rRNA sequencing. Depending on application purposes, both methods are suitable for identification of microbial communities, with FL-ONT being more superior at species level.

## 1. Introduction

The effect of tumour associated microbial communities on tumour biology is under intense investigation (1–4). To date, the tumour microbiome has been implicated in modulating anti-tumoural immune responses, chemotherapy efficacy, and tumour progression (2–4). Apart from tissues, microbial signatures from other collection sites such as stool and saliva may have diagnostic or prognostic roles in various cancers (1, 5, 6). Together these studies demonstrate the potential impact of understanding the tumour microbiome in cancers. However, as a prerequisite to further research, it is critical to use the right tools for a robust microbiome identification.

DNA sequencing techniques such as targeted sequencing of the 16S ribosomal RNA (rRNA) gene, metagenomics, and to a lesser extent, meta-transcriptomics have been instrumental in microbiome identification (4). Of these, Illumina based short-read sequencing (SRS) of the 16S rRNA has been widely adopted due to its relatively low cost and high throughput (4, 7). The 16S rRNA gene is approximately 1,500 to 1,600 base pairs (bp) long in most bacteria, and is composed of nine variable regions which allows taxonomical identification of microbial communities. Although sequencing all nine variable regions offers better taxonomic resolution, most studies usually sequence only a selection of variable regions, limiting the capacity of species level identification (8).

In head and neck cancer (HNC), most studies on microbiome identification relied on SRS of the 16S V3-V5 region on tissues, swabs, saliva, and oral rinse (8, 9). Our recent meta-analysis of V3-V5 short-read Illumina sequencing datasets identified key oral microbes localised in HNC tumours (8). However, taxonomic classifications were limited to the genus level, with species-specific contributions to HNC pathophysiology largely unknown (8, 10). Given that several oral species such as *Fusobacterium nucleatum* and *Porphyromonas gingivalis* can promote tumour progression and alter anti-tumour immunity (11), utilising cutting-edge technologies that can provide species level information will provide critical insights to the role of microbiome in HNC.

Long read sequencing (LRS) technologies from Oxford Nanopore Technologies (ONT) or Pacific Biosciences (PacBio) have been rapidly improving (ONT: Quality Score (Q-score) > 20, PacBio: Q-score > 33) and applied in the mainstream for various DNA sequencing applications, enabling sequencing of longer reads (> 10,000 bp) (12). Importantly, its application in full-length 16S rRNA gene sequencing enables in-depth taxonomic classification (12). Numerous studies have compared ONT LRS to Illumina based 16s rRNA gene sequencing in mock communities, swabs, and faecal samples (13–26). Four studies investigated the difference in beta-diversity (13, 18, 24, 26), with two studies showing differences in beta-diversities between ONT and Illumina (13, 18). Two other studies measured sum of agreement at genera level (sum of the percentage of matching genera) and showed that the median microbiome agreement between ONT and Illumina groups ranged from 65% to 70% (18, 26). Moreover, many studies have analysed the differences or correlations between the abundance estimates generated by ONT and Illumina sequencing technologies at different taxonomic levels (13–26). The consensus is that at higher taxonomic levels, there were greater correlation observed, while the least correlation was observed at species level (14, 19, 21, 26). To date, there has been no comparison between LRS and SRS using tumour tissue samples.

In this study, we comprehensively evaluated the differences in microbiome diversities and abundance between ONT and Illumina 16S rRNA sequencing technique on HNC tissue samples. Bacterial abundance between ONT and Illumina was evaluated at each taxonomic level using paired Wilcoxon test on relative abundance and paired ANOVA-Like Differential Expression tool 2 (ALDEx2) differential abundance analysis, which takes into account the compositional and zero-inflation nature of microbiome dataset (27). Furthermore, matrix assisted laser desorption ionization-time of flight mass spectrometry (MALDI-TOF MS) was also performed on bacteria isolated from 4 patient tissue samples for comparison to the 16S rRNA sequencing performed. To our best knowledge, this is the first study to perform long read 16S rRNA sequencing on HNC cancer tissue samples, and the first to evaluate ONT and Illumina 16S rRNA sequencing on HNC tissue samples.

## 2. Methods

### 2.1 Sample collections

Tumour samples were collected from 26 HNC patients undergoing surgical resection of primary tumours at the Royal Adelaide Hospital (Adelaide, SA, Australia) and The Memorial Hospital (Adelaide, SA, Australia). Tumour samples were placed into a sterile cryotube immediately after surgical excision to prevent any environmental contamination. Ethics approval for the collection and storage of patient samples were granted by Central Adelaide Local Health Network Human Research Ethics Committee (Adelaide, South Australia) (HREC MYIP14116), and all patients had signed written informed consent.

### 2.2 DNA extraction

DNA was extracted in a laminar flow cabinet with aseptic technique, using DNeasy Blood & Tissue Kit (Qiagen, Germany, Hilden) with some modification, as described previously (28). Briefly, prior to DNA extraction, the tissue samples were homogenised using 3 mm stainless steel beads (Qiagen) and a TissueLyser II (Qiagen) at 23 Hz for 3 minutes. Afterwards, the homogenized tissues were incubated with 1 mg/mL lysozyme (cat no: L3790, Sigma Aldrich, MO, USA) and 0.2 mg/mL lysostaphin (L7386, Sigma) at 37°C for 1 hour, followed by 0.5 mg/mL proteinase K (Qiagen) incubation at 56°C for 2 hours, before proceeding with manufacturer’s DNA extraction protocol. The DNA was quantified using Qubit™ dsDNA Quantification Assay Kit (Invitrogen, USA, MA), before undergoing Illumina 16S rRNA V3-V4 SRS (referred to as V3V4-Illumina) and ONT full-length V1-V9 16S rRNA LRS (referred to as FL-ONT). Negative controls were also included in extraction process.

### 2.3 V3V4-Illumina 16S rRNA sequencing

PCR amplification and sequencing was performed by the Australian Genome Research Facility (Adelaide, SA, Australia). PCR amplicons were generated using V3-V4 primers and conditions as described previously (26). Thermocycling was completed with an Applied Biosystem 384 Veriti and using Platinum SuperFi II master mix (Invitrogen) for the primary PCR. The first stage PCR was cleaned using magnetic beads, and samples were visualised on 2% SYBR E-gel (ThermoFisher). A secondary PCR to index the amplicons was performed with the same polymerase master mix. The resulting amplicons were cleaned again using magnetic beads, quantified by Quantifluor fluorometry (Promega, USA) and normalised. The equimolar pool was cleaned a final time using magnetic beads to concentrate the pool and then measured using a High-Sensitivity D1000 Tape on an Agilent 2200 TapeStation (Agilent Technologies, CA, USA). The pool was diluted to 5nM and molarity was confirmed again using a Qubit High Sensitivity dsDNA assay (ThermoFisher). This was followed by sequencing on an Illumina MiSeq (Illumina, CA, USA) with a V3, 600 cycle kit (2 x 300 bp paired-end).

### 2.4 FL-ONT 16S rRNA sequencing

Full-length V1-V9 sequencing was performed using ONT MinION workflows (Oxford Nanopore Technologies, Oxford, UK). Full length 16S rRNA were amplified using 16S Barcoding Kit (SQK-16S024, Oxford Nanopore Technologies), with PCR conditions described in Supplementary Table 2. Amplicons were purified using AMPure® XP beads (Beckman Coulter Diagnostics, USA, CA), quantified using Qubit HS kit (Qiagen), before sequencing on R9.4.1 chemistry (FLO-MIN106) flow cells (Oxford Nanopore Technologies), following manufacturer’s protocol. Basecalling was conducted using the super-accuracy basecalling model with Guppy v6.2.11.

### 2.5 Pre-processing and taxonomy assignment

For FL-ONT, EMU, was used to estimate full-length 16S rRNA relative abundance (10). For V3V4-Illumina, taxonomy assignment was performed using Divisive Amplicon Denoising Algorithm 2 (DADA2) (29). All taxonomic assignment were performed using SILVA reference database v11.5 (10, 29). Paired samples with low read counts (< 1000) after taxonomy alignment were removed. Negative controls were filtered at this step, as they had no read counts.

### 2.6 Alpha- and Beta-diversity analysis

Since short-read Illumina 16S rRNA sequencing is limited to genus level resolution (10), we performed alpha and beta-diversities analyses at the genus level. Data was agglomerated to genus level, and a total of 155 genera were identified. Alpha-diversity was measured using Shannon, Simpson, InvSimpson and Observed indexes for each sample were calculated using R package, microeco (30). Wilcoxon matched-pairs signed rank test was performed to determine differences between paired samples sequenced using different techniques.

For beta-diversity analysis, CLR-abundance (offset = 0.5) of all genera were ordinated using Euclidean distance and plotted on a principal coordinate analysis (PCoA) using phyloseq v1.46 and ggpubr v0.6 R packages (31). Permutational multivariate analysis of variance (PERMANOVA) and Analysis of similarities (ANOSIM), strata for paired sample, were performed to assess differences between in beta-diversity between paired ONT and Illumina sequencing groups (32). Additionally, we also included the W_d_ test, a test which is robust for heteroscedastic datasets, to determine differences in beta-diversity between ONT and Illumina (33). Variance between groups were measured using the betadisper test from vegan v2.6 (32). Permutations for all tests were set to n = 9999.

### 2.7 Differential abundance analysis

We analysed differential abundance at all taxonomic levels – phylum, class, order, and family, genus, and species. Data was agglomerated to specific levels before downstream analysis. For differential relative abundance analysis, data was normalized into relative abundance (%), and Wilcoxon matched-pairs signed rank test adjusted for false discovery rate (FDR) was used to determine differences between paired samples. Additionally, we also applied ALDEx2 differential abundance analysis, which uses a Monte Carlo Dirichlet sampling approach which considers the compositional and zero-inflation nature of microbiome dataset while to determining differences between ONT and Illumina sequencing group (27).

### 2.8 Culture-based identification

Additional homogenised tumour tissues from four patients (PT09, PT14, PT15, PT17) were also cultured on sheep blood agar plates (Thermofisher) in an anaerobic hood at 37°C condition, 0% O_2_, 5% CO_2_, 2.5 % H_2_. Bacteria isolates were sent for MALDI-TOF MS for identification.

## 3. Results

### 3.1 Study workflow and patient demographics

Tumour tissue samples from 26 HNC patients were collected for FL-ONT and V3V4-Illumina (Figure 1, Supplementary Table 1). Data were processed using DADA2 for Illumina or EMU for ONT data. The SILVA v11.5 rRNA database was used for taxonomy alignment for 16s rRNA data generated from both sequencing techniques (Figure 1). After sample processing and agglomerating to each taxonomic level, the total counts of unique phyla, classes, orders, families, genera, and species detected were as follows: 15 phyla, 20 classes, 52 orders, 89 families, 155 genera, and 225 species.

**Figure 1:**
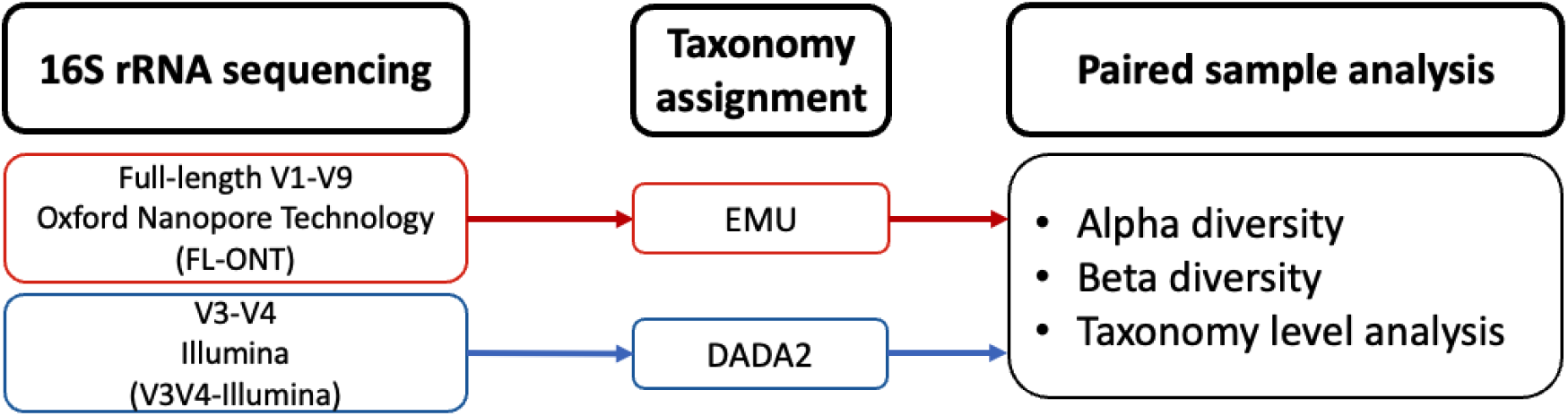
Workflow and data processing.

### 3.2 FL-ONT and V3V4-Illumina 16S rRNA sequencing groups display comparable alpha diversity indexes at the genus level

To compare observed richness and evenness between FL-ONT and V3V4-Illumina, alpha diversity was measured using Shannon, Simpson, InvSimpson, and Observed indexes (Figure 2A-2D). Since Illumina SRS 16S rRNA sequencing is largely limited to genus level resolution, alpha diversity was measured at genus level (10). After agglomerating datasets to genus level, a total of 155 genera were identified. Similar to previous findings comparing LRS and SRS (18), we found no significant differences (p < 0.05) between ONT and Illumina 16S rRNA sequencing – Shannon (mean difference = −0.230, 95% CI = −0.420 to −0.040), InvSimpson (mean difference = −0.765, 95% CI = −2.052 to 0.521) and Observed genuses (mean difference = 0.923, 95% CI = −2.910 to 4.756) (Figure 2A, 2C, 2D). However, Simpson index (mean difference = −0.07, 95% CI = −0.124 to −0.019, p =0.02) showed statistically significant but small differences between groups (Figure 2B). Overall, these results suggest that there are minimal differences between ONT and Illumina 16S rRNA sequencing groups with respect to alpha diversity at the genus level.

**Figure 2:**
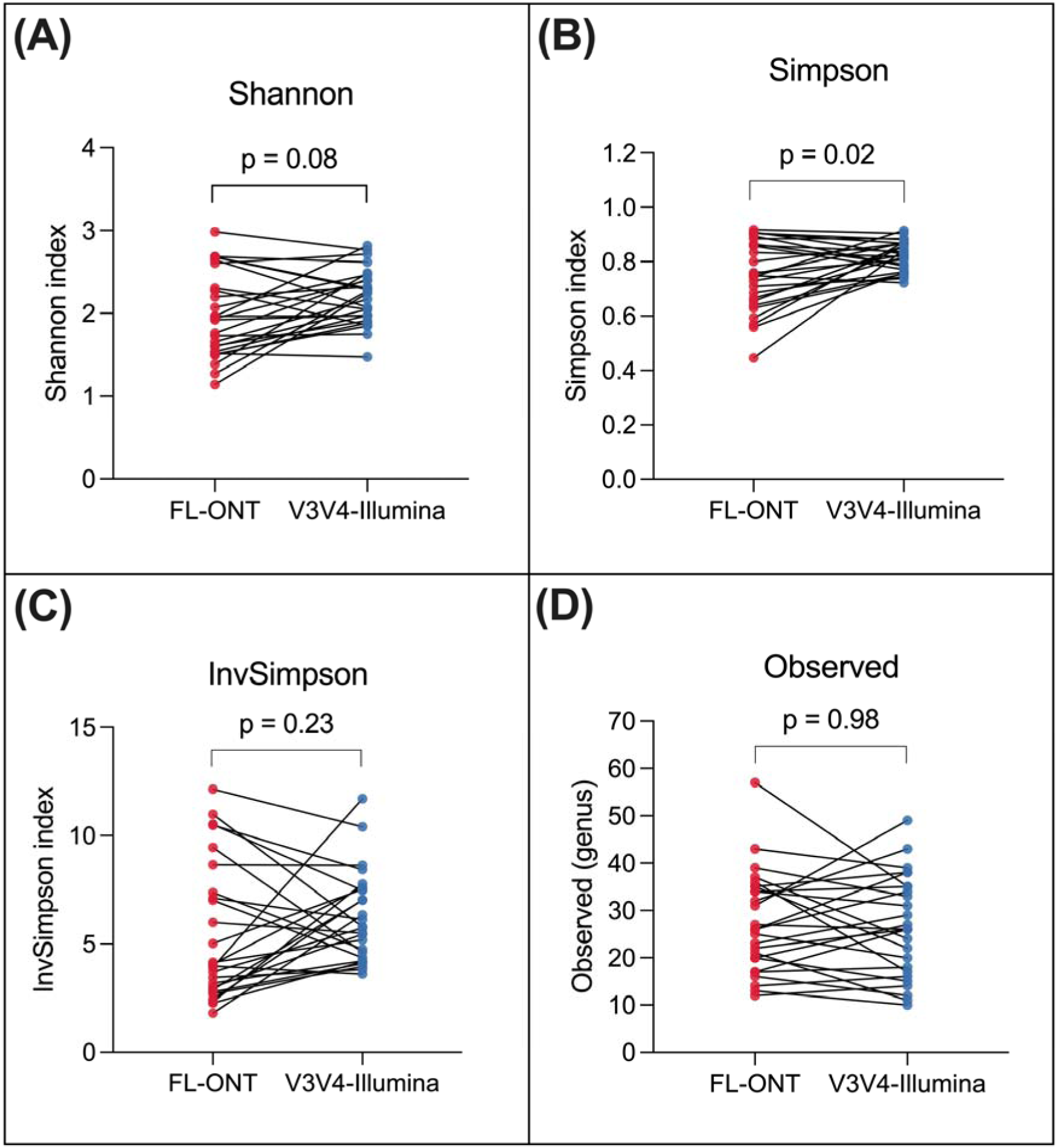
Paired alpha diversity analysis of FL-ONT and V3V4-Illumina at the genus level. Tissues were sequenced using ONT and Illumina technologies and data were aligned to the SILVA 16S rRNA database. To compare the differences in alpha diversity between technologies, paired Wilcoxon rank sum tests (adjusted for FDR) was performed for (A) Shannon index, (B) Simpson index, (C) InvSimpson, and (D) Observed features (genus) using R package, microeco.

### 3.3 Differences in beta-diversity were observed between paired FL-ONT and V3V4-Illumina sequencing on tumour samples at the genus level

Differences in β-diversity between FL-ONT and V3V4-Illumina were assessed using PCoA plot of Euclidean distance on CLR normalized abundance, PERMANOVA, ANOSIM and W_d_ test (Figure 3). Ordination PCoA Euclidean plot suggest that there is a shift in beta diversity between FL-ONT and V3V4-Illumina 16S rRNA sequencing (Figure 3). Similarly, we observed significant differences in β-diversity between FL-ONT and V3V4-Illumina using PERMANOVA test (PERMANOVA - R^2^ = 0.083, p < 0.0001). Dissimilarities between groups were assessed using an ANOSIM test (R = 0.25, p < 0.0001), further showing significant differences between both sample groups (Figure 3). However, significant differences in dispersion were observed between both technologies (Permutest – p < 0.001, F = 0.0035). Given that ANOSIM, and to a lesser extent, PERMANOVA, can be influenced by differences in dispersion across groups (34), we have additionally incorporated W_d_ test, a test which is robust for heteroscedastic datasets (33). Similar to PERMANOVA and ANOSIM, W_d_ test also showed significant differences between in β-diversity between FL-ONT and V3V4-Illumina 16S rRNA sequencing (W_d_ = 4.63, p = 0.0001) (Figure 3). Taken together, these findings show that β-diversity differs between FL-ONT and V3V4-Illumina 16S rRNA sequencing at the genus level.

**Figure 3:**
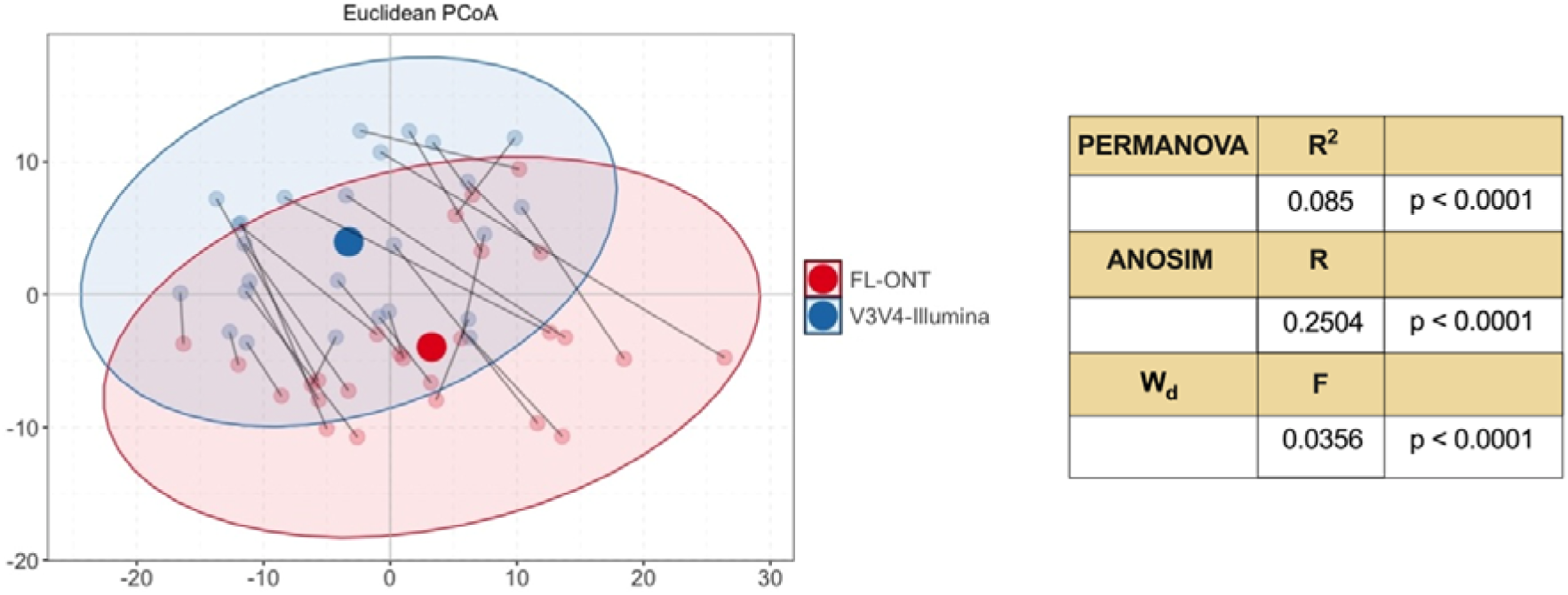
Paired beta diversity analysis of paired FL-ONT and V3V4-Illumina 16S rRNA sequencing on tissue samples at the genus level. Principal Coordination Analysis (PCoA) plot of Euclidean distance on CLR normalized abundance. PERMANOVA, ANOSIM and W_d_ test were performed, statistics and p-value were presented. Red and blue dot-points represents ONT and Illumina 16S rRNA sequencing respectively, while line between dot-points represents paired samples.

### 3.4 Paired sample analysis of FL-ONT and V3V4-Illumina 16S rRNA sequencing reveals differences at higher taxonomic levels - phylum, class, order, and family

To determine taxonomic differences at phylum, class, order, and family level between FL-ONT and V3V4-Illumina sequencing technologies, we performed paired Wilcoxon rank sum test on CLR-normalized abundance using ALDEx2 (Figure 4, Supplementary Table S3-S6, Supplementary Figure S2-S3) (27).

**Figure 4:**
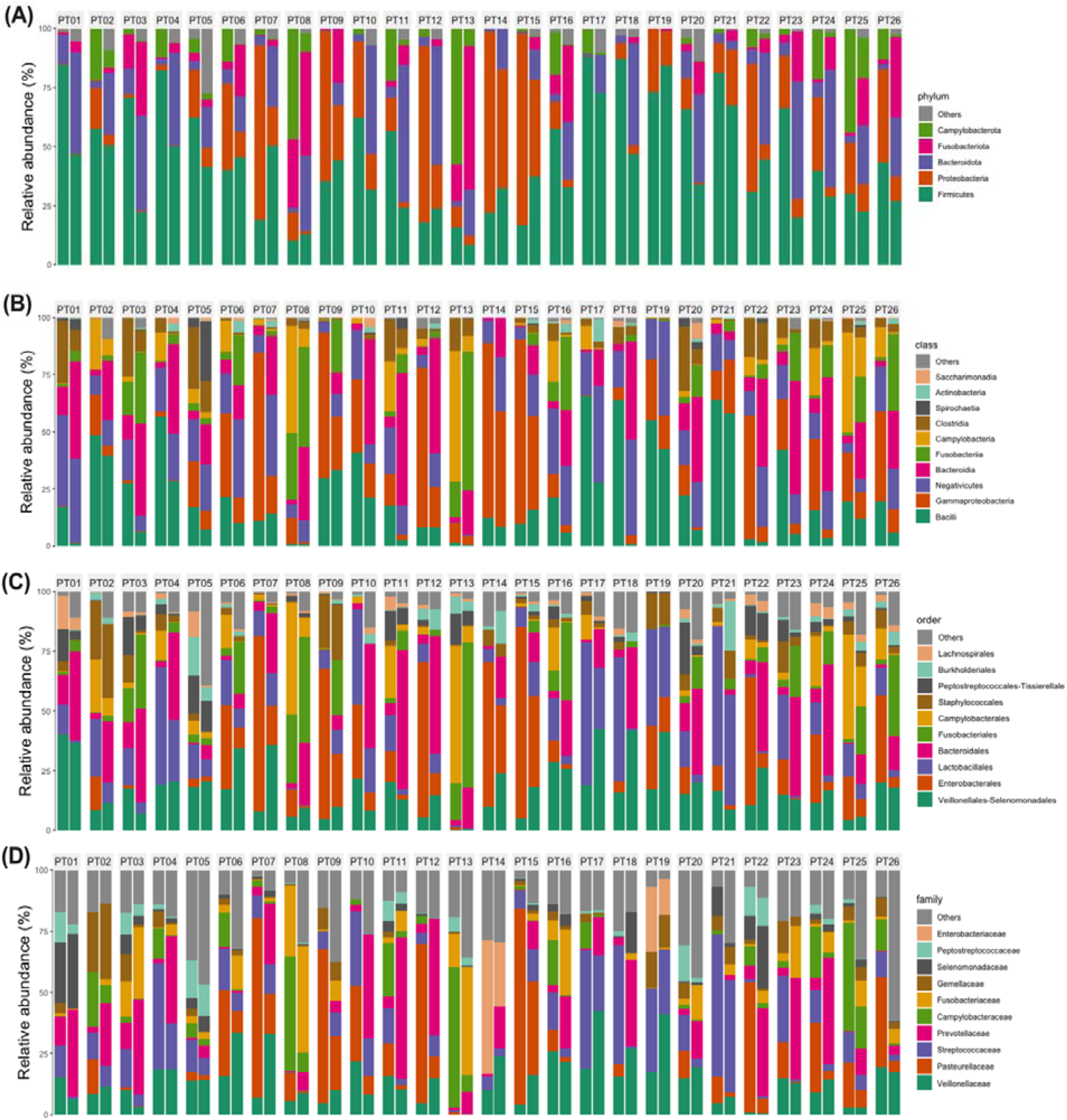
Relative abundance comparison between FL-ONT and V3V4-Illumina 16S rRNA sequencing from phylum to family levels. Relative abundance of top (A) Phylum, (B) Class, (C) Order, and (D) Family after agglomerating to each level. For each patient panel (PT01-PT26), the FL-ONT is shown on the left and V3V4-Illumina on the right. Paired Wilcoxon tests were performed to compare differences between FL-ONT to V3V4-Illumina sequencing (Supplementary Table S3-S6).

#### Phylum level

Based on relative abundance, the main phylum detected in both techniques were *Campylobacterota* (FL-ONT: 12.07%, V3V4-Illumina: 2.07%)*, Fusobacteriota* (FL-ONT: 3.50%, V3V4-Illumina: 13.95%)*, Bacteroidota* (FL-ONT: 4.14%, V3V4-Illumina: 29.39%)*, Proteobacteria* (FL-ONT: 28.98%, V3V4-Illumina: 10.90%)*, Firmicutes* (FL-ONT: 50.63%, V3V4-Illumina: 38.56%) (Figure 4A, Table S3A). *Proteobacteria* (mean diff = 18.08%, p < 0.0001)*, Firmicutes* (mean diff = 12.08%, p < 0.01)*, and Campylobacterota* (mean diff = 10.01%, p < 0.0001) were significantly more abundant in FL-ONT group, while *Fusobacteriota* (mean diff = −10.45%, p < 0.0001) and *Bacteroidota* (mean diff = −25.24%, p < 0.0001) were more abundant in V3V4-Illumina group (Figure 4A, Figure S2A, Table S3A). Overall, FL-ONT and V3V4-Illumina sequencing showed a moderate correlation (R > 0.67) between groups at the phylum level (mean R = 0.8402) (Table S3B, Figure S1). Furthermore, among top 5 phyla detected in FL-ONT, 4/5 phyla were also top phyla detected in V3V4-Illumina (Table S3A). Additionally, we applied ALDEx2 differential analysis and identified significantly (p < 0.05) lower CLR-abundance of *Campylobacterota* (CLR_diff.btw_ = −5.270, effect size = −2.424)*, Proteobacteria* (CLR_diff.btw_ = −4.066, effect size = −2.802)*, Firmicutes* (CLR_diff.btw_ = −2.580, effect size = −1.674), and higher CLR-abundance in *Actinobacteriota* (CLR_diff.btw_ = 2.946, effect size = 0.781) in V3V4-Illumina, as compared to FL-ONT (Figure S3B, Table S3C).

#### Class level

Based on relative abundance, we found that 12/20 bacterial classes were significantly different among sequencing groups, and 5 of these classes had a have mean difference of more than 10% (Figure 4B, Figure S2B, Table S4A). The FL-ONT group had greater abundance of *Gammaproteobacteria* (mean diff = 17.99%, p < 0.0001)*, Bacilli* (mean diff = 13.23%, p < 0.0001)*, and Campylobacteria* (mean diff = 10.01%, p < 0.0001), while *Bacteroidia* (mean diff = −25.24%, p < 0.0001) and *Fusobacteriia* (mean diff = −10.45%, p < 0.0001) were greater in V3V4-Illumina group (Figure 4B, Figure S2B, Table S4A). Overall, FL-ONT and V3V4-Illumina sequencing showed moderate correlation (R > 0.65) between groups at the class level (Mean R = 0.8305) (Table S4B, Figure S1). Moreover, among top 10 classes detected in FL-ONT, 8/10 classes were also among the top classes detected in V3V4-Illumina (Table S4A). Using ALDEx2, we found 6/20 bacterial classes that were significantly different (p < 0.05) between sequencing technologies (Figure S3B, Table S4C). *Campylobacteria* (CLR_diff.btw_ = −5.151, effect size = −2.565)*, Gammaproteobacteria* (CLR_diff.btw_ = −4.001, effect size = −2.773), *Bacilli* (CLR_diff.btw_ = −3.369, effect size = −2.137)*, Clostridia* (CLR_diff.btw_ = −2.726, effect size = −1.619)*, Negativicutes* (CLR_diff.btw_ = −1.538, effect size = −1.019) were significantly lower CLR-abundance in V3V4-Illumina group, while V3V4-Illumina group were determined to contain more *Actinobacteria* (CLR_diff.btw_ = 3.112, effect size = 0.749) (Figure S3B, Table S4C). Notably, all six bacterial classes were lineage to Phylum *Campylobacterota, Proteobacteria, Actinobacteriota and Firmicutes* (Table S3-S4). Similarly, there were bacterial classes that were only detected in FL-ONT or V3V4-Illumina groups, albeit being < 1% mean relative abundance (Table S4A).

#### Order level

When comparing relative abundance at the order level, we identified 14/52 orders being significantly different between FL-ONT and V3V4-Illumina 16S rRNA sequencing group (Figure 4C, Figure S2C, Table S5A). *Enterobacterales* (mean diff = 18.88%, p < 0.0001)*, Lactobacillales* (mean diff = 11.19%, p < 0.0001)*, Campylobacterales* (mean diff = 10.01%, p < 0.0001) were significantly higher in FL-ONT groups, while *Fusobacteriales* (mean diff = - 10.45%, p < 0.0001) and *Bacteroidales* (mean diff = −22.34%, p < 0.0001) were higher in V3V4-Illumina sample group (Figure S2C, Table S5A). Overall, the correlation between FL-ONT and V3V4-Illumina at the order level (mean R = 0.6460) lower than phylum and class levels (Table S5B, Figure S1). Among the top 10 bacteria order detected in FL-ONT, 9/10 were also top order detected in V3V4-Illumina (Table S5A). Using ALDEx2 differential abundance analysis, we identified 9/52 bacterial orders that were significantly different (p < 0.05) between sequencing technologies (Figure S3C, Table S5C). *Xanthomonadales* (CLR_diff.btw_ = - 7.824, effect size = −2.703)*, Campylobacterales* (CLR_diff.btw_ = −4.986, effect size = −2.254)*, Enterobacterales* (CLR_diff.btw_ = −4.536, effect size = −2.385)*, Lactobacillales* (CLR_diff.btw_ = −3.370, effect size = −1.768)*, Staphylococcales* (CLR_diff.btw_ = −3.221, effect size = −1.678)*, Peptostreptococcales-Tissierellales* (CLR_diff.btw_ = −2.745, effect size = −1.408)*, Lachnospirales* (CLR_diff.btw_ = −2.081, effect size = −1.126), and *Veillonellales-Selenomonadales* (CLR_diff.btw_ = −1.375, effect size = −0.868) have lower CLR-abundance in V3V4-Illumina samples, while *Micrococcales* (CLR_diff.btw_ = 5.802, effect size = 1.211) is more abundant in V3V4-Illumina group (Figure S3C, Table S5C). All these orders were lineages of classes that were significantly different between sequencing technique groups (Table S4-S5). However, orders *Xanthomonadales* and *Micrococcales* were the only detected in FL-ONT and V3V4-Illumina sequencing group respectively.

#### Family level

When comparing differences in relative abundance between FL-ONT and V3V4-Illumina groups, we identified differences in 23/89 families (Figure 4D, Table S6A). Of these families, *Pasteurellaceae* (mean diff = 17.22%, p < 0.0001), and *Campylobacteraceae* (mean diff = 10.01%, p < 0.0001) were significantly more abundant in FL-ONT groups, while *Prevotellaceae* (mean diff = −19.97%, p < 0.0001) was significantly more abundant in V3V4-Illumina groups (Figure S2D, Table S6A). Overall, the correlation between FL-ONT and V3V4-Illumina at the family level (mean R = 0.5205) were lower than all the higher taxonomic levels (Table S6B, Figure S1). Among the top 10 bacteria families detected in FL-ONT, 7/10 were also top families detected with V3V4-Illumina (Table S6A). Using ALDEx2 differential abundance analysis, 10/89 bacterial families were significantly different (p < 0.05) between sequencing technologies. The V3V4-Illumina group exhibited lower CLR-abundance of *Streptococcaceae* (CLR_diff.btw_ = −3.193, effect size = −1.689)*, Campylobacteraceae* (CLR_diff.btw_ = −4.889, effect size = −2.142)*, Xanthomonadaceae* (CLR_diff.btw_ = −7.849, effect size = −2.689)*, Carnobacteriaceae* (CLR_diff.btw_ = −3.282, effect size = −0.7724)*, Veillonellaceae* (CLR_diff.btw_ = −1.702, effect size = −0.9191)*, Lachnospiraceae* (CLR_diff.btw_ = −1.865, effect size = −0.9609)*, Pasteurellaceae* (CLR_diff.btw_ = −4.757, effect size = −2.189) and *Gemellaceae* (CLR_diff.btw_ = −3.143, effect size = −1.574), while contain more *Burkholderiaceae* (CLR_diff.btw_ = 4.689, effect size = 0.9494) and *Micrococcaceae* (CLR_diff.btw_ = 5.570, effect size = 1.011), as compared to the FL-ONT group (Figure S3D, Table S6C). Importantly, *Burkholderiaceae* and *Micrococcaceae* were only detected by V3V4-Illumina group (Table S6A). *Burkholderiaceae* was the only family that does not come from bacterial order that were significantly different in our ALDEx2 analysis (Table S5B, S6B).

Overall, the bacteria identified by FL-ONT and V3V4-Illumina group were mostly from the same lineage at the phylum, class, order, and family taxonomical levels. However, we also detected bacteria that were unique to the sequencing technique, albeit detected at very low abundance (< 0.1%) (Table S3-S6). Furthermore, we observed decreasing correlation between the relative abundance of FL-ONT and V3V4-Illumina group from higher (phylum) to lower (family) taxonomic groups (Figure S1). Finally, we also observed that there was a good concordance in the relative abundance of the top bacteria detected, whereby both techniques have similar top bacteria detected.

### 3.5 FL-ONT and V3V4-Illumina 16S rRNA sequencing displays greater discrepancies in microbial community profiling at the genus level

Since Illumina 16S rRNA SRS is capable of identifying taxa mostly to the genus level, with limited capability of identification at species level, we compared FL-ONT and V3V4-Illumina at the genus level (10, 35). When we compared relative abundance between sequencing techniques, we found that 29/155 bacterial genera were significantly different in relative abundance (Figure 5A-5B, Table S7A). *Haemophilus* (mean diff = 15.02%, p < 0.0001) and *Campylobacter* (mean diff = 10.05%, p < 0.0001) had significantly higher relative abundance in FL-ONT group, while *Prevotella_7* (mean diff = −12.04 %, p < 0.0001) had significantly higher relative abundance in the V3V4-Illumina group (Figure 5B, Table S7A). Other notable bacterial genera such as *Streptococcus* (mean diff = 9.575%, p < 0.0001) and *Fusobacterium* (mean diff = −6.816%, p = 0.00002) also had significantly higher relative abundances in FL-ONT and V3V4-Illumina group respectively (Table S7A). Of these 29 bacterial genera, 22 bacterial genera had less than 5% differences in relative abundance between techniques, although being statistically significant (Table S7A). Notably, a moderate correlation (Mean R = 0.5205) between FL-ONT and V3V4-Illumina was observed at the genus level (mean R = 0.5205) (Table S7B, Figure S1). Among the top 10 genera detected in FL-ONT, only 6/10 genera were among the top genera detected in V3V4-Illumina (Table S7A).

**Figure 5:**
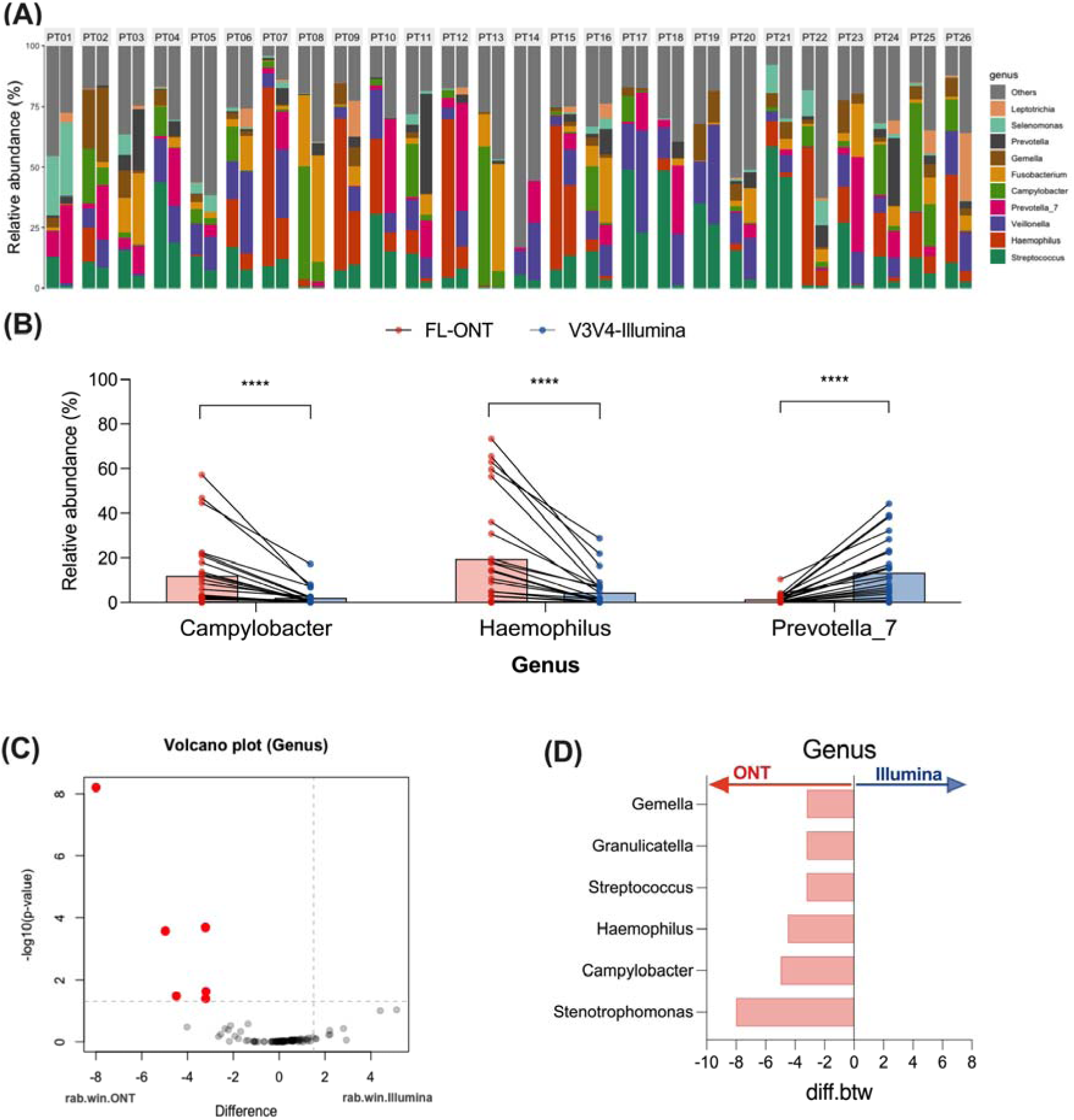
Comparison of abundance between FL-ONT and V3V4-Illumina 16S rRNA sequencing at the Genus level. After agglomerating to genus level, a total of 155 genera were identified. (A) Relative abundance (%) of top 10 genus, strata to per patient. For each patient panel, ONT and Illumina sequencing were represented by left and right bar plot respectively. (B) Relative abundance (%) of genus with > 10% differences between techniques. Paired Wilcoxon tests were performed to compare differences between ONT to Illumina sequencing. Additionally, ALDEx2 was performed to assess differences in genus between sequencing techniques. (C) ALDEx2 volcano plot. Red dot points represent Benjamini-Hochberg corrected p-value of Wilcoxon test < 0.05. Rab.win.group refers to the median bacterial clr value for the group of samples. (D) Genera that were significantly different between ONT and Illumina using ALDEx2 analysis. Diff.btw refers to the median difference in bacterial clr values between ONT and Illumina groups (Illumina - ONT). ****p < 0.0001

Using ALDEx2 differential abundance test, 6/155 bacterial genera were significantly different in CLR-abundance between the sequencing technologies (Figure 5C-5D, Table S7C). All 6 bacterial genera were lower in CLR-abundance in Illumina samples as compared to ONT samples (Figure 5C-5D, Table S7C) – *Streptococcus* (CLR_diff.btw_ = −3.215, effect size = −1.616)*, Campylobacter* (CLR_diff.btw_ = −4.973, effect size = −2.073)*, Stenotrophomonas* (CLR_diff.btw_ = −7.998, effect size = −2.876)*, Granulicatella* (CLR_diff.btw_ = −3.207, effect size = −0.732)*, Haemophilus* (CLR_diff.btw_ = −4.493, effect size = −1.662), and *Gemella* (CLR_diff.btw_ = −3.195, effect size = −1.626). Regardless of sequencing technique, these genera were detected in all samples. Notably, the family of these 6 genera were also significantly different when comparing FL-ONT to V3V4-Illumina 16S rRNA sequencing (Table S6C).

### 3.6 ONT LRS Full-length 16S rRNA sequencing is superior for species level bacterial identification

Illumina SRS is limited to sequencing short fragments which results in poor capacity to differentiate and identify highly similar species (10, 35). By sequencing the full-length 16S rRNA gene, FL-ONT is able to provide bacterial community identification at the species level. We further compared FL-ONT to V3V4-Illumina in HNC tissues samples at the species level. Furthermore, we also isolated bacteria from 4 HNC patients and identified these bacteria using MALDI-TOF MS to confirm that FL-ONT were able to identify the correct bacterial species.

A total of 225 bacteria species were identified among both sequencing groups (Table S8A). Of these 225 bacterial species detected, 96 (42.7%) were identified by both sequencing approaches. We detected 88/225 (39.1%) bacterial species that were unique to the FL-ONT group, with 79/88 of these bacterial species containing less than 1% of abundance (Figure 6A, Table S8A). For V3V4-Illumina sequencing, 41/225 (18.2%) species were unique to the group and critically all 41 of these bacterial species were less than 1% abundance in V3V4-Illumina group (Figure 6A, Table S8A).

**Figure 6:**
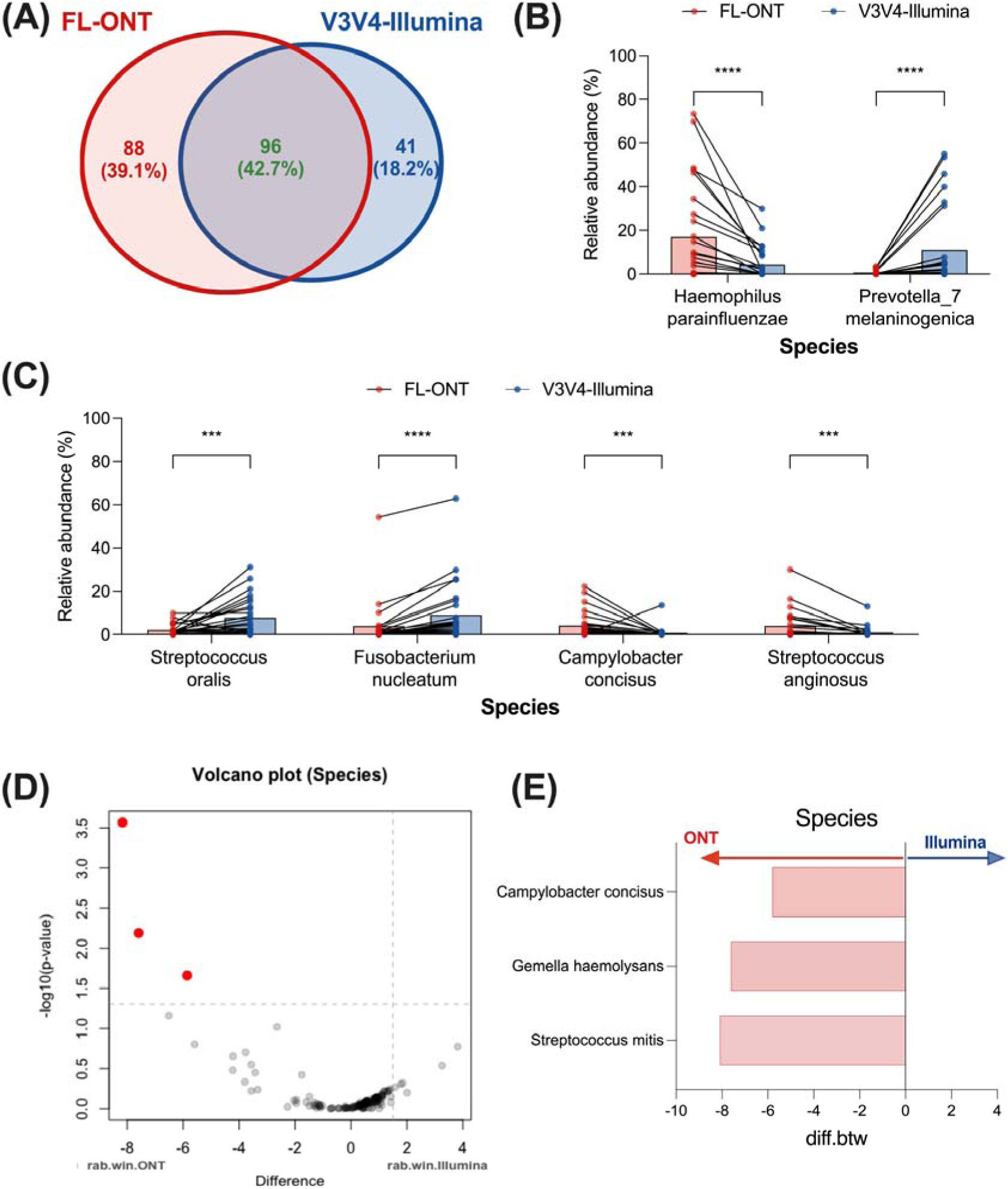
Comparison of abundance between FL-ONT and V3V4-Illumina 16S rRNA sequencing at the species level. After agglomerating to species level, a total of 225 species were identified. (A) Venn diagram of identified species in FL-ONT and V3V4-Illumina. Identified species is defined as having > 0 % abundant (Table S9A). (B) Relative abundance (%) of significantly different species with mean differences > 10% between ONT and Illumina groups. (C) Relative abundance (%) of significantly different species with mean differences > 3% between FL-ONT and V3V4-Illumina. Paired Wilcoxon test was performed to compare differences between FL-ONT to V3V4-Illumina sequencing. Additionally, ALDEx2 was performed to assess differences in species between sequencing techniques. (D) ALDEx2 volcano plot. Red dot points represent Benjamini-Hochberg corrected p-value of Wilcoxon test < 0.05. Rab.win.group refers to median bacterial species clr value for the group of samples. (E) Species that were significantly different using ALDEx2 analysis. Diff.btw refers to median difference in bacterial species clr values between FL-ONT and V3V4-Illumina groups (Illumina - ONT). ****p < 0.0001, ***p < 0.001.

Based on relative abundance, we identified 20 bacterial species to be significantly different (p < 0.05) between sequencing groups (Figure 6B-6C, Table S8A). Comparing relative abundance of FL-ONT to V3V4-Illumina, *Haemophilus parainfluenzae* (mean diff = 12.55%, p < 0.001) is more abundant in FL-ONT, while *Prevotella_7 melaninogenica* (mean diff = −10.41, p < 0.001) is more abundant in V3V4-Illumina sequencing, with both having a >10% differences in relative abundance (Figure 6B). Additionally, other species with substantial differences (> 3%) between FL-ONT and V3V4-Illumina groups includes *Campylobacter concisus* (mean diff = 3.365%, p < 0.01)*, Streptococcus anginosus* (mean diff = 3.018%, p < 0.05%)*, Streptococcus oralis* (mean diff = −5.692, p = 0.0107), and *Fusobacterium nucleatum* (mean diff = −5.174, p = 0.00136) (Figure 6C, Table S8B). Furthermore, 6/20 of these bacterial species (*Gemella haemolysans, Peptostreptococcus stomatis, Gemella sanguinis, Streptococcus salivarius, Streptococcus pseudopneumoniae, Lachnoanaerobaculum orale*) were not found in V3V4-Illumina group, while only *Rothia mucilaginosa* was not detected by FL-ONT group (Table S8A). As expected, we observed a poor correlation between FL-ONT and V3V4-Illumina at the species level (mean R = 0.2395) (Table S8B, Figure S1). Furthermore, FL-ONT and V3V4-Illumina only shared 4/10 of the top 10 detected species (Table S8A). Using ALDEx2 differential abundance analysis, similar findings were shown where the CLR-abundance of *Streptococcus mitis* (CLR_diff.btw_ = −8.097, effect size = −1.746), *Gemella haemolysans* (CLR_diff.btw_ = −7.616, effect size = −1.266) and *Campylobacter concisus* (CLR_diff.btw_ = −5.804, effect size = −1.223) were significantly lower in V3V4-Illumina, as compared to FL-ONT group (Figure 6D-6E, Table S8C), confirming differential abundance analysis. Furthermore, MALDI-TOF MS identified 16 isolates from four patients (Table S9). Overall, 81.3% of total isolates identified by MALDI-TOF MS were also identified using FL-ONT, while V3V4-Illumina was able to only identify 62.5% of isolates (Table S9A).

As expected, we observed large discrepancies in bacterial genus and species identification between FL-ONT and V3V4-Illumina sequencing groups. Moreover, FL-ONT was able to identify more unique bacterial species at a higher bacterial abundance than V3V4-Illumina sequencing. All the species unique to V3V4-Illumina were less than 1% relative abundance. Furthermore, MALDI-TOF MS identification were more identical to FL-ONT than V3V4-Illumina. Similar to previous studies, poor correlation between FL-ONT and V3V4-Illumina was observed at genus and species level (Figure S1) (14, 19, 21).

## 4. Discussion

Recent studies suggest that microbiome contributions to tumour pathobiology can be attributed to specific bacterial species (2–4), there is a significant need need to adopt sequencing technologies capable of species level identification such as FL-ONT 16S rRNA sequencing (10). We have previously reported a consensus tissue microbiome signature in HNC using previously published Illumina SRS 16S rRNA sequencing data (8). To the best of our knowledge, this is the first study to perform FL-ONT 16S rRNA sequencing on HNC tumour samples. Furthermore, we comprehensively assessed the performance of FL-ONT to V3V4-Illumina sequencing. We found that alpha diversity was comparable between paired FL-ONT and V3V4-Illumina 16S rRNA sequencing. In contrast, beta-diversity was significantly different between paired FL-ONT and V3V4-Illumina 16S rRNA sequenced HNC samples. At higher taxonomic levels (phylum, class, order, and family), moderate correlations between the two sequencing methodologies for bacterial relative abundance, while at lower taxonomic levels, including genus and species, the correlations were poor correlation. Importantly, FL-ONT identified more unique species that were also detected at higher in abundance than V3V4-Illumina.

In this study, we compared alpha and beta-diversities between FL-ONT to V3V4-Illumina 16S rRNA sequencing data at the genus level which is the current acceptable limit for short-read Illumina 16S rRNA sequencing based taxonomic classification (10). Similar to our previous work on nasal swabs (26), we identified comparable alpha-diversities between paired FL-ONT and V3V4-Illumina 16S rRNA sequencing in HNC tissues samples. Out of the 4 alpha diversities matrices tested, only Simpson index showed a statistically significant, but minimal difference (mean differences = −0.07) in our study. A previous study also reported minimal but statistically significant differences in alpha-diversity measurement using InvSimpson index between the two sequencing techniques (18). Importantly, in our HNC tissue samples, we showed minimal or no differences in alpha-diversities. Consistent with our findings, previous reports have shown significant beta-diversity differences between ONT and Illumina based 16S rRNA sequencing in the gut and nasal microbiome (13, 18). Critically, our beta-diversities were stratified for patients accounting for inter-patient sample differences. Together, these findings indicate that ONT and Illumina 16S rRNA sequencing have minimal impact on bacterial genera richness and evenness, however overall bacterial composition was affected by the sequencing technique employed.

We next determined whether bacterial composition difference observed were present in every taxonomic level. Previous studies have examined FL-ONT and Illumina 16S rRNA sequencing datasets for differences in relative abundance at the phylum (13), order (14), family (14–16, 26), genus (14–22, 24–26), and species (14, 16, 19, 20, 26) level. However, these studies have used different analytical approaches that may affect the interpretation of their results. Some compared relative abundance of paired samples without paired differential abundance analysis (13, 22, 24), while others compared averages within each sequencing group (18, 19). Most compared correlation in abundance between ONT and Illumina (14, 19, 21, 25), specifically the top 10 to 15 bacteria (14, 15, 19, 21), thus not reflecting the magnitude of differences in abundance between sequencing techniques. Furthermore, a few studies had small sample sizes (< 10) which limits their interpretation (13, 14, 17, 22). Notably, in addition to this study, our previous study on nasal swabs was the only study to have applied paired analysis to evaluate differences in relative abundance (family, genus) and diversities between ONT and Illumina sequencing (26). Paired differential abundance analysis should be employed to account for inter-sample differences such as lifestyle activities including smoking, alcohol or diet intake that is known to affect the microbiome (36–38).

Consistent with most studies (14, 19, 21), we observed a decrease in correlation between microbial abundance produced from different sequencing techniques from higher to lower taxonomic levels (Figure S1). In our study, we observed differences in relative abundance between FL-ONT and V3V4-Illumina 16S rRNA sequencing especially for bacteria related to phylum *Campylobacterota, Proteobacteria, Actinobacteriota and Firmicutes*. Furthermore, we found that there were biases in the bacteria detected in FL-ONT or V3V4-Illumina 16S rRNA sequencing. ALDEx2 is an alternative method that considers compositional and zero-inflated microbiome datasets and is more robust than standard relative abundance analyses (39). Using ALDEx2, we also identified differences at every taxonomy although at a smaller number, reflective of its conservative nature to reduce false-postives detection (39, 40). Taken together, we have comprehensively shown that there are significant differences in the two sequencing technologies’ ability to detect the microbial composition of HNC tissues.

The microbiome has been reported to influence numerous facets of tumour pathobiology biology including treatment efficacy, tumour immunity and tumour progression (1). Gemcitabine, a chemotherapeutic treatment for pancreatic, bladder and metastatic triple-negative breast cancers, can be transported into the cytoplasm of *Gammaproteobacteria* (class) using nucleoside transporter (NupC), where it gets inactivated by bacterial cytidine deaminase (41–43). Gut-derived *Bifidobacterium* spp. is associated with increased response rates and progression free survival to PD-1 checkpoint inhibitors (44). Notably, well-studied microbial metabolites such as butyrate, can also improve PD-1 checkpoint inhibitor response rates (45, 46). Butyrate can be produced from *Faecalibacterium* (genus) and *Akkermansia muciniphila* (species) (45, 46). Of note, these tumour modulating abilities is dependent on specific genomic features shared within a taxonomic level (43). Thus, microbiome identification at higher taxonomical levels that can be accurately identified by Illumina 16S rRNA sequencing is important (7). However, our study shows that FL-ONT 16S rRNA sequencing is similar to the precision of V3V4-Illumina at higher taxonimical levels but with the advantage of providing species level identification in a cost-effective manner. Using bacterial isolate cell culture and MALDI-TOF MS identification, we also showed that FL-ONT was able to identify more bacterial isolates compared to V3V4-Illumina sequencing. Thus, at higher taxonomic levels, Illumina 16S rRNA remains a cost-effective and accurate method to screen for microbial community (7, 47). However, with continued advances in the development of LRS technologies full-length 16S rRNA sequencing is quickly becoming a more attractive alternative with its ability for specied level microbial community classification (7, 47).

Although we have thoroughly investigated differences in both techniques, there are limitations to this study. Firstly, this study did not include an oral mock microbial community as a reference. Having a commercial oral mock community will allow benchmarking of library preparation steps such as primer efficacy and PCR conditions between both FL-ONT and V3V4-Illumina 16S rRNA sequencing. Furthermore, future studies should consider including other primers or all primer sets to cover the entire region of the 16S rRNA for short-read Illumina sequencing. This will ensure better coverage and comparison between full-length ONT and full-length short-read Illumina sequencing (48). Additionally, future studies should also include more samples and culture conditions (i.e. aerobic and anaerobic) in the culturomics approach. Lastly, adding on a metagenomics approach can also provide greater confidence with extra sequencing coverage outside of the 16S rRNA gene (7).

In conclusion, our study provides the first comprehensive comparison of FL-ONT and V3V4-Illumina 16S rRNA microbial sequencing in HNC tumour tissue samples. We have shown that there were key differences such as beta-diversity and some bacterial groups in every taxonomy at every level. Critically, we show that FL-ONT can provide more information about the microbiome that is cost-effective. We expect this technology to be more widely adopted in future cancer microbiome studies.

## Data Availability Statement

All sequencing data in this paper will be provided upon request.

## Author Contributions

Conceptualization, K.Y. and K.F.; methodology, investigation and data analysis, K.Y., J.C., E.S., G.B. W.M., R.V.; resources, J.H, S.K, A.P., P.W. and S.V.; writing – original draft preparation, K.Y., E.S., S.V. and K.F; writing – review and editing, K.Y., J.C., W.M., G.B., E.S., A.P., P.W., R.V., S.V. and K.F.; supervision, R.V., A.P., S.V. and K.F.; funding acquisition, A.P., P.W. and S.V. All authors have read and agreed to the published version of the manuscript.

## Acknowledgments

This work is supported by an NHMRC investigator grant APP1196832 to P.W., a The Garnett Passe and Rodney Williams Senior Fellowship to S.V., and The University of Adelaide Postgraduate Research Scholarship to K.Y. We would like to thank the medical staff from The Royal Adelaide Hospital and The Memorial Hospital for their assistance in sample collection.

## Ethics statement

Ethics approval for the collection and storage of patient samples were granted by Central Adelaide Local Health Network Human Research Ethics Committee (Adelaide, South Australia) (HREC MYIP14116), and all patients had signed written informed consent.

