## Supplementary Figures for "A comparison between full-length 16S rRNA Oxford Nanopore sequencing and Illumina V3-V4 16S rRNA sequencing in head and neck cancer tissues"

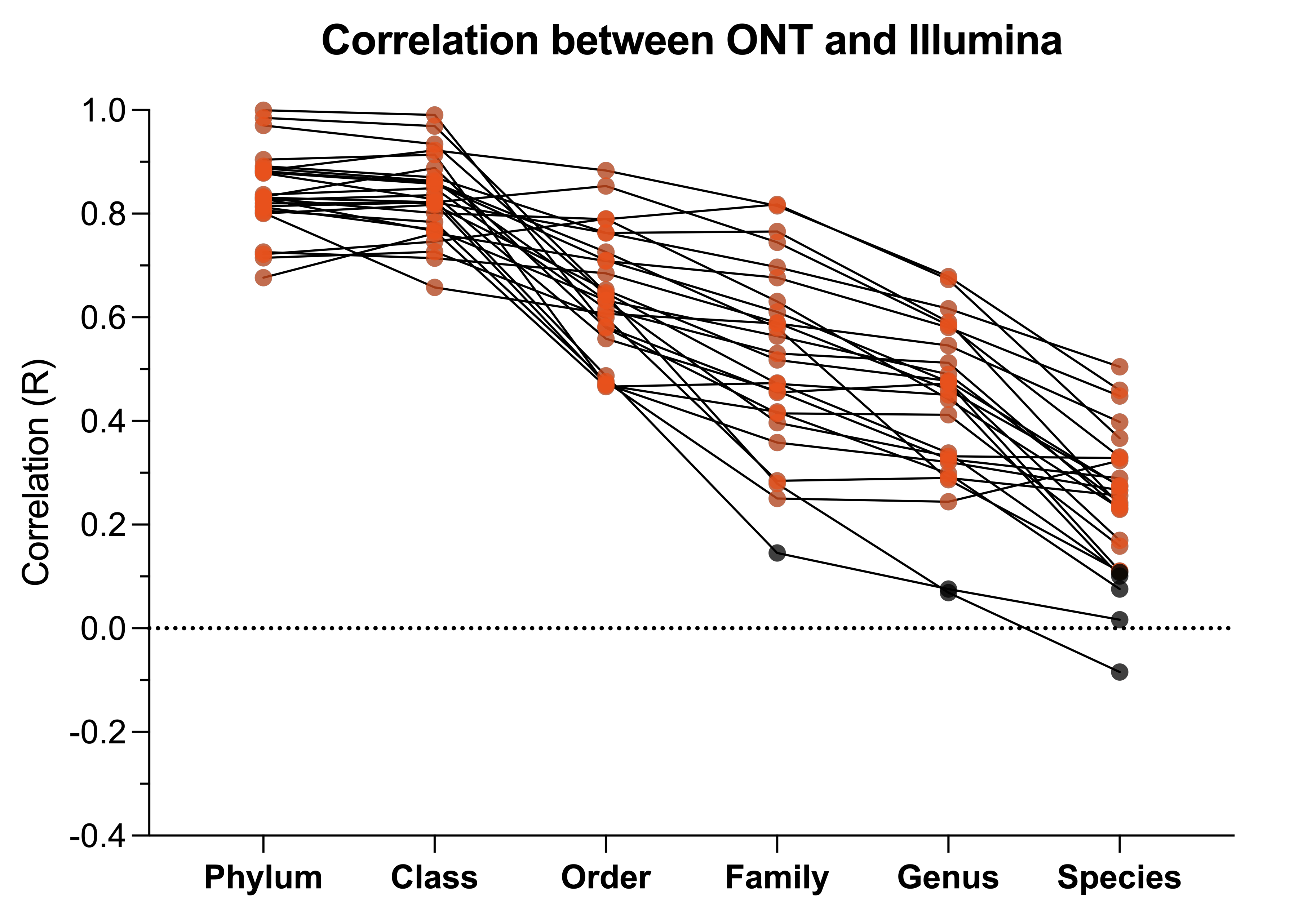

**Figure S1: Correlation in bacterial relative abundance (%) at every taxonomic level between FL-ONT and V3V4-Illumina 16S rRNA sequencing.** Spearman **c**orrelation analysis was performed for paired FL-ONT and V3V4-Illumina groups at each taxonomy level. Each point represents correlation value (R) between FL-ONT and V3V4-Illumina in each patient, and red points represent p < 0.05.

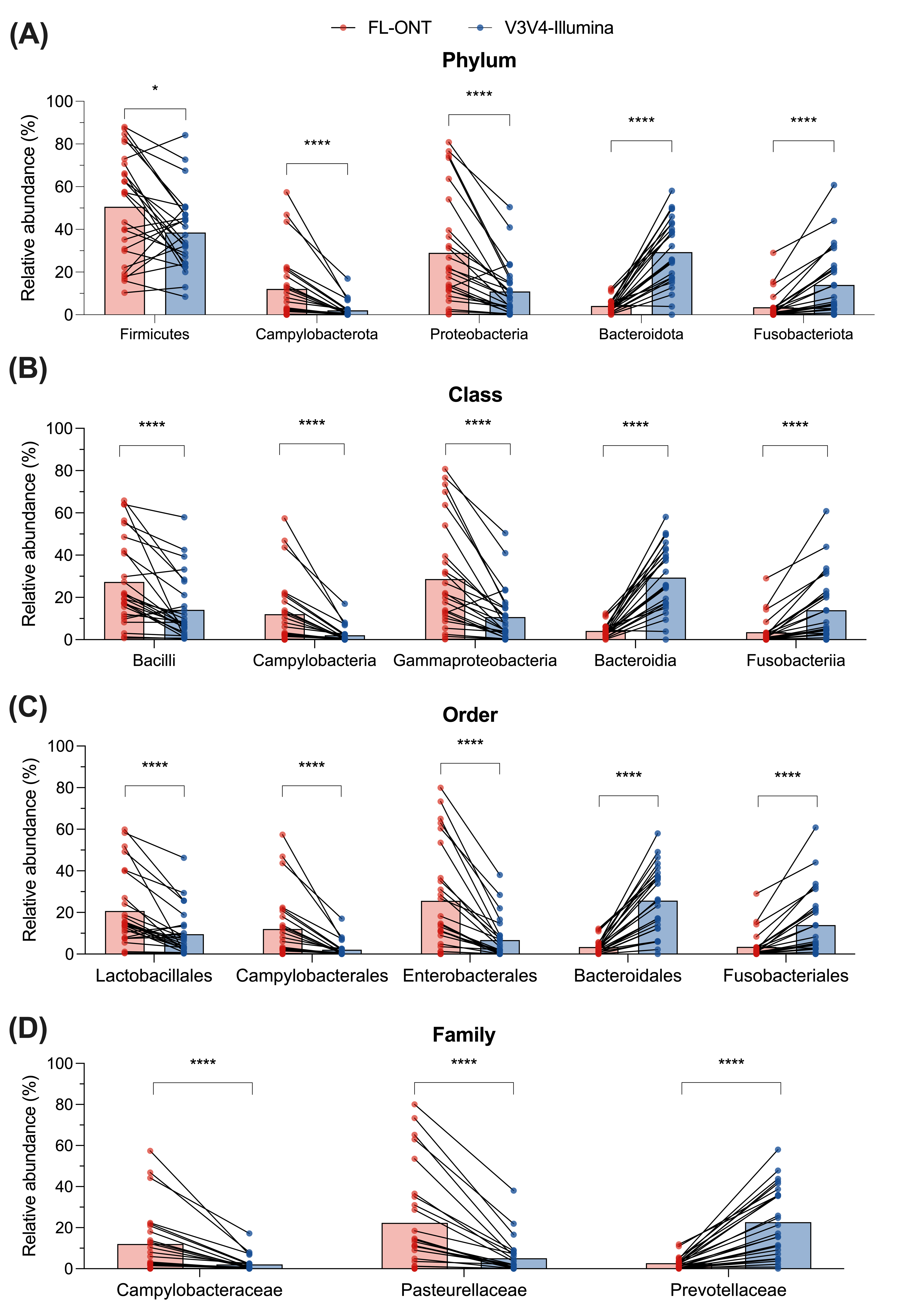

**Figure S2: Comparison of relative abundance between FL-ONT and V3V4-Illumina 16S rRNA sequencing at (A) Phylum, (B), Class, (C) Order and (D) Family levels.** After agglomerating to each taxonomy level, relative abundance was compared between both sequencing techniques (Supplementary Table S3-S6). Relative abundance (%) at each taxonomic levels with > mean 10% differences between techniques. Paired Wilcoxon were performed to compare differences between ONT to Illumina sequencing. ****p < 0.0001, *p < 0.05.

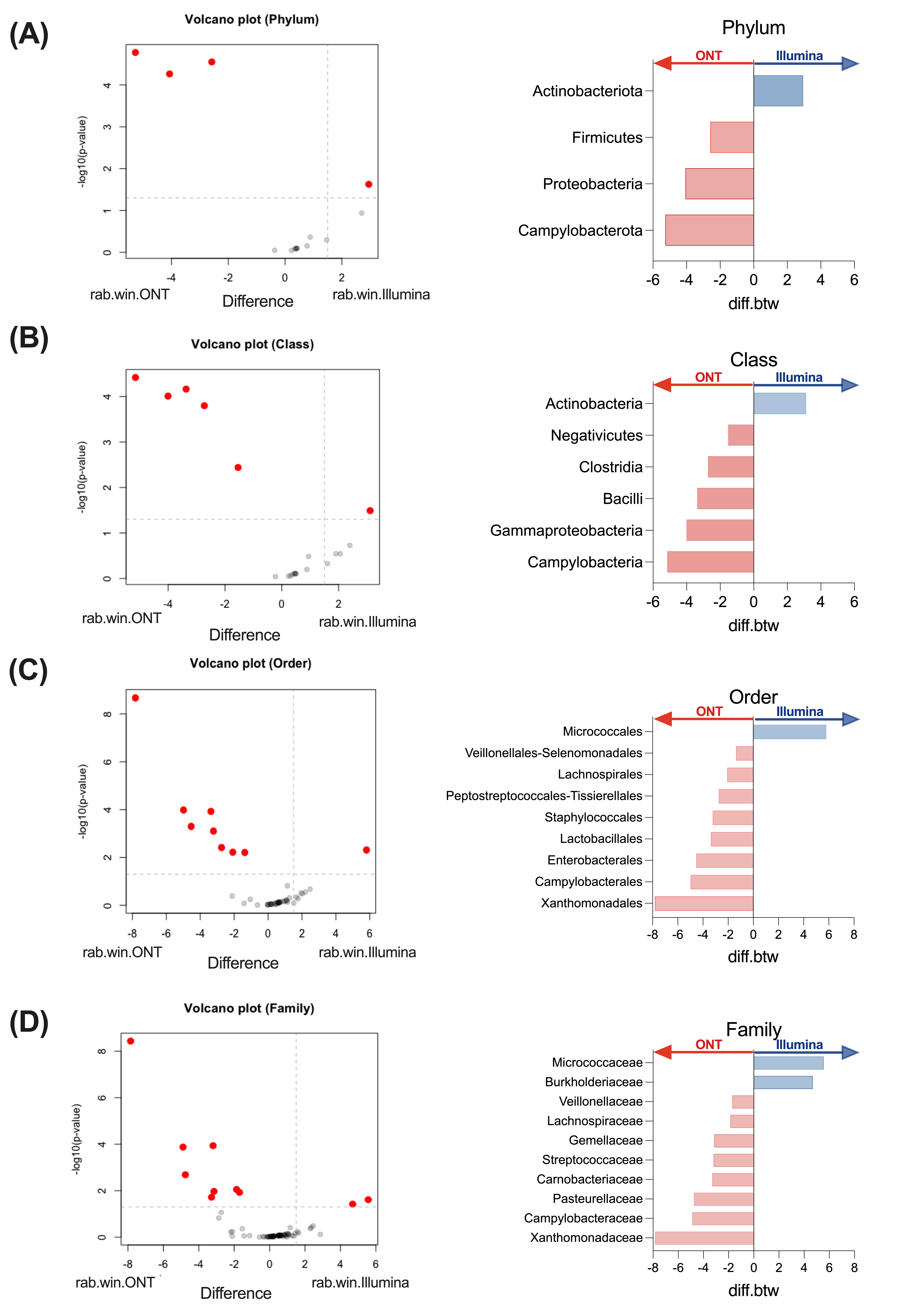

**Figure S3: ALDEx2 analysis show taxonomic differences between FL-ONT and V3V4-Illumina 16S rRNA sequencing.** ALDEx2 analysis was applied at (A) Phylum, (B) Class, (C) Order, and (D) Family levels. Volcano plot (left panel) shows differences in bacteria from phylum to family levels between ONT and Illumina. Red points represent -log_10_(p-value) < 0.05. Points to the left of 0 represent enrichment in Illumina, to the right represent enrichment in ONT. Taxa that were significantly different at all levels were represented in bar plots (right panel). The of bar plot represents bacteria being more abundant in FL-ONT (red) or V3V4-Illumina (blue) sequencing technique.
